# Structural basis for the initiation of COPII vesicle biogenesis

**DOI:** 10.1101/2020.10.08.331793

**Authors:** Aaron M.N. Joiner, J. Christopher Fromme

## Abstract

The first stage of the eukaryotic secretory pathway is the packaging of cargo proteins into COPII vesicles exiting the endoplasmic reticulum (ER). The cytoplasmic COPII vesicle coat machinery is recruited to the ER membrane by the activated, GTP-bound, form of the conserved Sar1 GTPase. Activation of Sar1 on the surface of the ER by Sec12, a membrane-anchored GEF (guanine nucleotide exchange factor), is therefore the initiating step of the secretory pathway. Here we report the structure of the complex between Sar1 and the cytoplasmic GEF domain of Sec12, both from *Saccharomyces cerevisiae*. This structure, representing the key nucleotide-free activation intermediate, reveals how the potassium ion-binding K-loop disrupts the nucleotide binding site of Sar1. We describe an unexpected orientation of the GEF domain relative to the membrane surface and propose a mechanism for how Sec12 facilitates membrane insertion of the amphipathic helix exposed by Sar1 upon GTP-binding.

## Introduction

The eukaryotic secretory pathway is essential for protein secretion and trafficking of virtually all proteins that localize to the plasma membrane, lysosomes, endosomes, and the Golgi complex. The first step of the secretory pathway, after protein synthesis and quality control measures within the ER, is the concentration and packaging of cargo into vesicles for transport to the Golgi complex. These vesicles are generated by the conserved coat protein complex II (COPII) machinery that interacts directly with the cytoplasmic portions of cargos and sculpts the ER membrane into transport carriers (Russell and Stagg, 2010; Jensen and Schekman, 2011; Zanetti et al., 2012; Barlowe and Miller, 2013; Barlowe, 2019).

The generation of COPII-coated vesicles at the ER is initiated by activation of Sar1, a small Ras-like GTPase of the Arf family (Oka et al., 1991; Barlowe et al., 1993; Huang et al., 2001; Bi et al., 2002). Activated, GTP-bound Sar1 triggers formation of the COPII coat by directly interacting with the inner layer of the coat complex, the Sec23/Sec24 heterodimer. Sar1 binds to the ER membrane through its N-terminal amphipathic helix, which is exposed only upon GTP-binding and plays a primary role in membrane remodeling during vesicle formation and fission (Antonny et al., 1997; Aridor et al., 2001; Huang et al., 2001; Bielli et al., 2005; Lee et al., 2005; Hariri et al., 2014; Schwieger et al., 2017).

Sec12 is the GEF responsible for activating Sar1 via nucleotide exchange (Nakano et al., 1988; d’Enfert et al., 1991; Barlowe and Schekman, 1993; Weissman et al., 2001; Futai et al., 2004). Among established GEFs for Arf family members, Sec12 is the only known integral membrane protein. Sec12 is anchored to the ER through its C-terminal transmembrane domain and the nucleotide exchange activity resides in the N-terminal cytosolic β-propeller domain (Barlowe and Schekman, 1993; Futai et al., 2004; McMahon et al., 2012).

Although the structures of Sec12 and Sar1 are known, and the kinetics of Sec12 GEF activity towards Sar1 have been characterized, the actual mechanism used by Sec12 to activate Sar1 via nucleotide exchange has not been determined (Huang et al., 2001; Bi et al., 2002; Futai et al., 2004; Rao et al., 2006; McMahon et al., 2012). In contrast to many small GTPases of the Ras superfamily, Sar1 and most other Arf GTPase family members are distinguished by the direct coupling of activation to insertion into the cytoplasmic leaflet of a membrane bilayer (Antonny et al., 1997). Despite extensive structural characterization of Arf family GEFs (Cherfils et al., 1998; Goldberg, 1998; Renault et al., 2003; DiNitto et al., 2007; Aizel et al., 2013; Malaby et al., 2013; Padovani et al., 2014; Galindo et al., 2016; Richardson et al., 2016; Karandur et al., 2017; Das et al., 2019), it has remained unresolved how these GEFs couple nucleotide exchange with membrane insertion of their substrates.

Here, we report the crystal structure of the yeast Sar1-Sec12 complex representing the key intermediate of the nucleotide exchange reaction. This structure provides detailed insights into the mechanism of nucleotide exchange, and in particular explains the vital role of the potassium ion-binding K-loop of Sec12 in rearranging the Sar1 nucleotide binding site. We have identified an extensive, positively charged surface on the complex which also contains the Sec12 transmembrane domain. The coexistence of these features on the same surface suggests a specific orientation of the complex on the membrane. We therefore propose that Sec12 positions Sar1 in an orientation that allows its N-terminal helix to insert directly into the ER membrane upon GTP binding.

## Results

### The structure of the Sar1-Sec12 nucleotide exchange intermediate

GEFs activate their GTPase substrates by competing with nucleotide for binding to the GTPase. Therefore, GEFs typically make extensive contacts with their substrate in order to displace bound GDP or GTP. As this binding is reversible and cellular GTP concentrations are much higher than GDP concentrations, subsequently the GEF is displaced by GTP resulting in activated GTPase (Goody and Hofmann-Goody, 2002). Nucleotide-free GTPase bound to its GEF is a critical intermediate in this pathway. To determine the structural and mechanistic basis for Sec12 GEF activity, we sought to capture the nucleotide-free intermediate of the Sar1-Sec12 guanine nucleotide exchange reaction. We incubated the purified cytosolic domain of *S. cerevisiae* Sec12 (residues 1-354) with a purified *S. cerevisiae* Sar1 construct lacking its N-terminal membrane-inserting amphipathic helix (residues 24-190) and a small amount of alkaline phosphatase. The phosphatase was included to hydrolyze guanine nucleotide which would otherwise compete with Sec12 for binding to Sar1. We purified the stable nucleotide-free Sar1-Sec12 complex (Figure S1) and grew crystals for X-ray diffraction studies. We collected an X-ray diffraction dataset using a single crystal and solved the structure by molecular replacement using the previously reported structures of yeast Sec12 (McMahon et al., 2012) and hamster Sar1-GDP (Huang et al., 2001). After model building and refinement against data to 2.3Å resolution (Figure S2), the final atomic model (Figure 1A) exhibits appropriate geometry and refinement statistics in light of moderate anisotropy of the diffraction data (Table 1).

**Figure 1:**
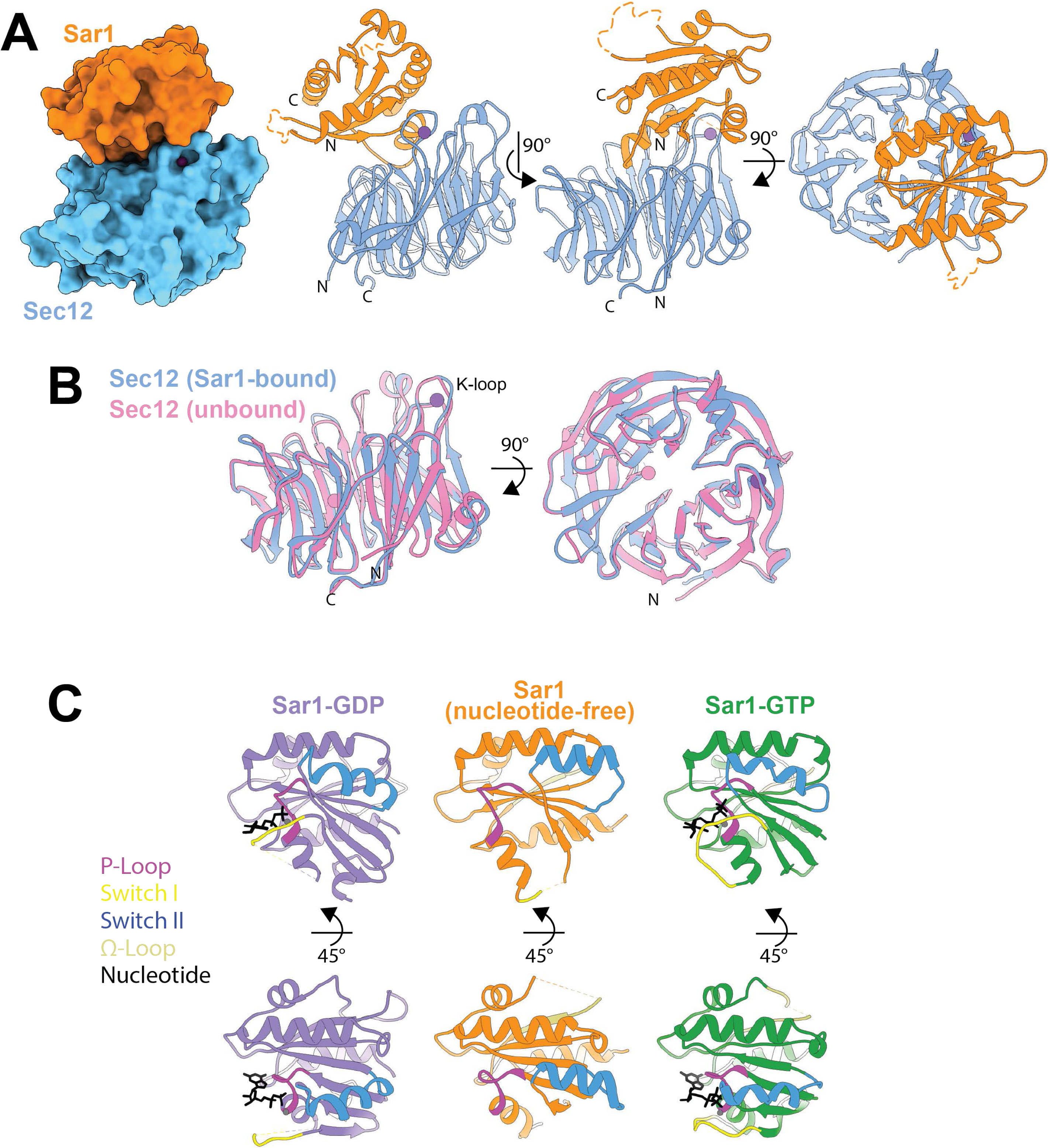
The structure of the Sar1-Sec12 nucleotide exchange intermediate. A. Sar1 (residues 24-190) is orange and Sec12 (residues 1-344) is blue. The purple sphere depicts the bound K^+^ ion in the K-loop of Sec12. Left, surface representation; right, ribbon depictions. The N- and C-termini are labeled and dashed lines indicate regions lacking electron density which were unmodeled. B. Superposition of Sar1-bound Sec12 (blue, this study), and Sec12 alone (pink, (McMahon et al., 2012), PDB: 4H5I). The pink sphere depicts a K^+^ ion that is observed only in the unbound crystal structure. C. Comparison of Sar1-GDP (purple, (Huang et al., 2001), PDB: 1F6B), nucleotide-free Sar1 (orange, this study), and Sar1-GTP (green, (Bi et al., 2002), PDB: 1M2O). Regions of Sar1 are highlighted by coloring: the phosphate loop (residues 30-37) is magenta, the switch I region (residues 46-57) is yellow, the switch II region (residues 76-92) is blue, the Ωloop (residues 154-169) is tan, and the nucleotide is black.

The structure of Sec12 in the Sar1-Sec12 complex is almost identical to the previously published structure of Sec12 alone (McMahon et al., 2012) (Figure 1B). Sec12 consists of seven β-sheets, or “blades”, arranged in a circular fashion to form a β-propeller. On one side of the first propeller blade, there is an extended loop, termed the K-loop, which binds to a potassium ion. This structural feature was previously demonstrated to be essential for GEF activity (McMahon et al., 2012). Consistent with these findings, we observe the K-loop makes direct contact with Sar1, an interface that is discussed further below. In addition, the conformation of the K-loop is essentially unchanged by Sar1-binding, indicating that it is a rigid structural element. In both the Sar1-bound and unbound structures, electron density is absent for the last ten residues of Sec12 (residues 345-354), suggesting these residues serve as a flexible linker connecting the catalytic cytosolic domain to the transmembrane domain (McMahon et al., 2012).

Sar1, like other small GTPases, adopts different structural conformations in its inactive (GDP-bound) and active (GTP-bound) forms (Huang et al., 2001; Bi et al., 2002; Rao et al., 2006; Bi et al., 2007). When bound to Sec12 as a nucleotide-free intermediate in the exchange reaction, we observed that Sar1 adopts a conformation significantly different from either its GDP- or GTP-bound forms (Figure 1C). The most significant observable differences between the structures correspond to the “switch” and “interswitch” regions, whereas the conformation of the phosphate loop remains largely unaltered by Sec12 binding. Some regions of Sar1 lack electron density in complex with Sec12, indicative of their flexibility: residues 48-65, which encompass the entirety of the switch I region and part of β3; and residues 152-162, which correspond to the Sar1-specific “Ωloop” (Huang et al., 2001), are disordered when bound to Sec12. The switch II region remains ordered but has shifted significantly due to displacement by the K-loop of Sec12. The switch II region moves by ~8 Å between GDP-bound and the Sec12-bound state and by ~10 Å from the Sec12-bound state to the GTP-bound state (Figure 1C). For comparison, the switch II region shifts ~8 Å between the GDP- and GTP-bound states (Figure 1C). The consequence of these structural changes is the loss of the nucleotide binding site, particularly the portion responsible for interacting with the Mg^2+^ ion required to stably bind to the negatively charged phosphate groups.

### The Sar1-Sec12 interface and the mechanism of nucleotide release

In the crystal structure, Sar1 is bound on one face of the Sec12 β-propeller, making contact with a surface that extends from the center of the β-propeller to the K-loop at the far edge of the first propeller blade (Figure 2A,B). This interface is extensive (1,040A^2^, Table S2) and displays a high degree of sequence conservation (Figure 2C).

**Figure 2:**
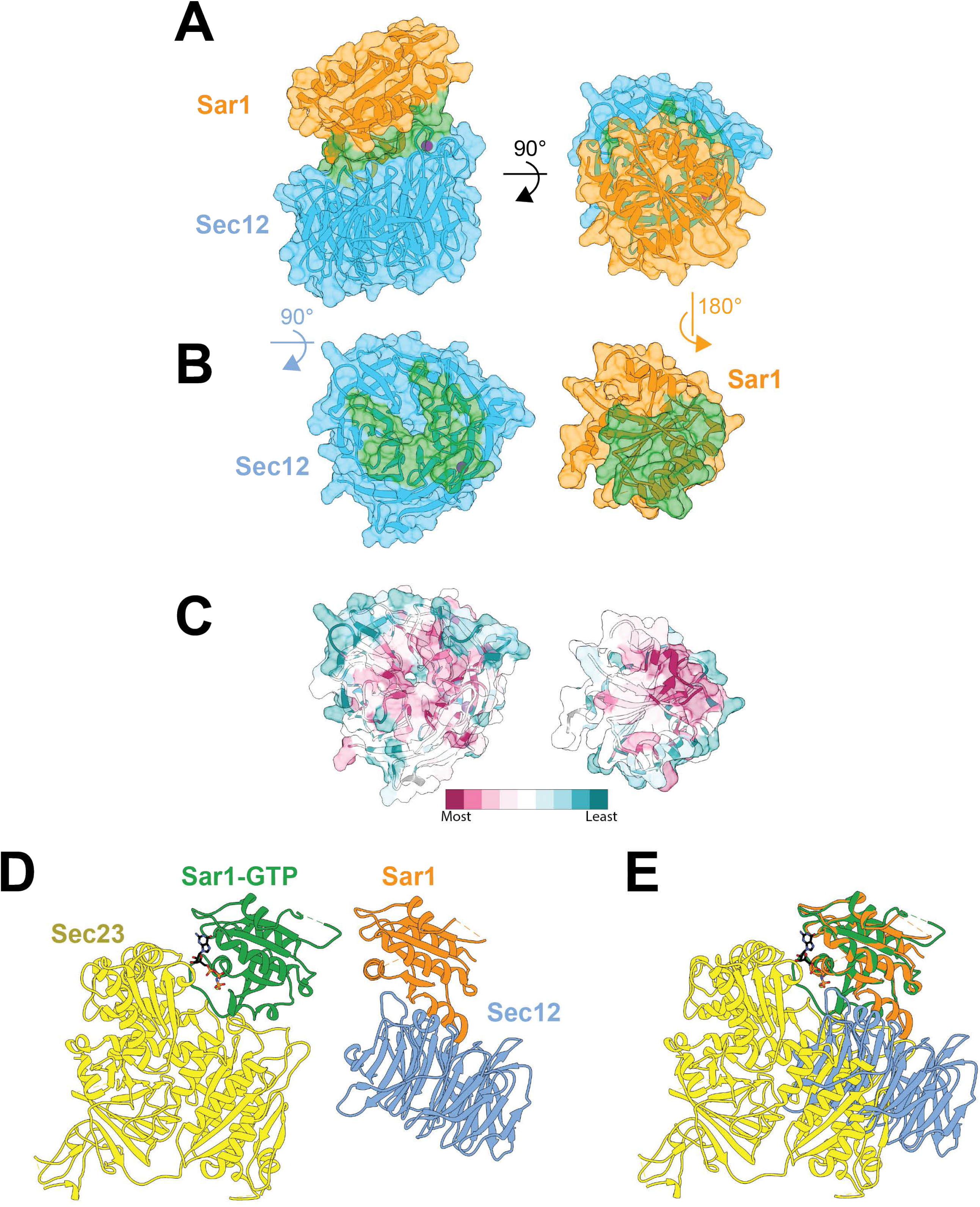
Sar1-Sec12 interaction surfaces. A. Transparent surface and ribbon representation of the Sar1-Sec12 complex in which the interfacial surface is colored green. The purple sphere depicts the bound K^+^ ion in the K-loop of Sec12. B. The complex (colored as in A) has been pulled apart in order to visualize the interface area (green) on each protein separately. C. Same as in B, but colored based on sequence conservation with maroon indicating highest conservation and teal representing low conservation. D. Ribbon representation of Sar1-GTP (green) bound to Sec23 (yellow) (Bi et al., 2002) next to nucleotide-free Sar1 (orange) bound to Sec12 (blue) (this study). E. As in D, but with Sar1 of each model superimposed.

There is significant overlap between the surface that Sar1 uses to bind to Sec12 and the surface that Sar1 uses to bind to its primary effector, Sec23 (Bi et al., 2002) (Figure 2D,E). As a consequence, we expect that Sar1-GTP bound to Sec23 in the COPII “pre-budding” complex will be protected from engaging with Sec12 again and undergoing unnecessary nucleotide exchange. Therefore, although GEFs do not exhibit a preference for acting upon GDP-bound versus GTP-bound GTPases *in vitro* (Goody and Hofmann-Goody, 2002), competition with Sec23 may effectively prevent Sec12 from non-productive nucleotide exchange of Sar1-GTP *in vivo*.

Previous studies investigated the functional importance of several conserved Sec12 residues (Futai et al., 2004; McMahon et al., 2012) (Table S3) that we now establish to lie at the Sar1 binding interface. The Sar1-Sec12 interface involves over 25 residues of Sar1, most of which belong to the switch II region, and over 30 residues of Sec12 from 6 out of the 7 blades of the β-propeller (Figure 3A). There are multiple types of interactions between residues of the two proteins (Figure S3). For example: hydrogen bonds are formed between the sidechains of Sec12 Tyr132 and Sar1 Lys85 and Asp86, and between the mainchain carbonyl oxygen of Sec12 Asn35 and the Sar1 Asp32 sidechain (Figures 3A,B and S3); an aromatic Pi-stacking interaction is observed between Sec12 Phe310 and Sar1 Phe72 (Figures 3A,B and S3); and an electrostatic interaction is observed between Sec12 Asp71 and Sar1 Arg82 (Figure 3C).

**Figure 3:**
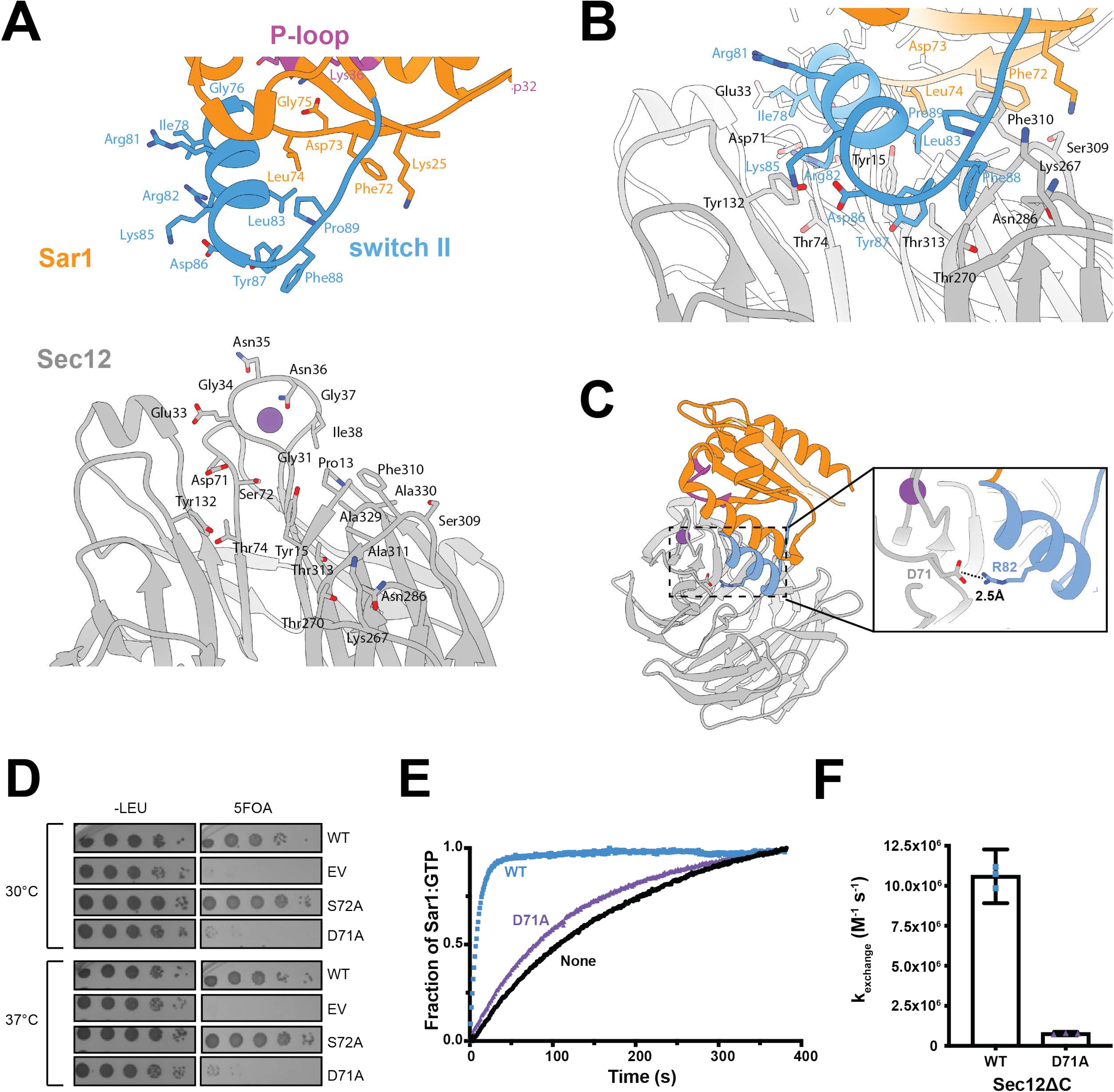
Sec12 interacts extensively with the Sar1 switch II region. A. Closeup view of the Sec12 and Sar1 interface with the molecules pulled apart to more clearly see the residues involved. The switch II region (blue) of Sar1 contributes many residues to the interaction surface with Sec12 (gray). Residues of the Sec12 K-loop and the surrounding propeller blades comprise the interaction surface. The purple sphere depicts the bound K^+^ ion in the K-loop of Sec12, and the Sar1 P-loop is pink. B. The interface viewed from a slightly different perspective. C. A broader perspective of the Sar1-Sec12 interface with zoomed inset highlighting the interaction between Sec12 Asp71 and Sar1 Arg82. D. The *sec12-D71A* mutant is unable to complement the loss of wild-type *SEC12*. E. Representative, normalized traces from *in vitro* nucleotide exchange assay measuring the kinetics of Sec12ΔC activation of full-length Sar1 in the presence of synthetic liposomes. F. Quantification of E. Wild-type exchange rate is similar to the previously reported rate (McMahon et al., 2012). Error bars indicate the 95% confidence intervals, n = 3.

While prior investigations tested the importance of some of these Sec12 residues, we identified many whose functional significance had not been interrogated. We generated several new mutant Sec12 alleles and screened them for their ability to complement a *sec12Δ* yeast strain (Figure 3D and Table S3). We observed that the *sec12-D71A* mutant was inviable (Figure 3D), suggesting that the electrostatic interaction between Sec12 Asp71 and Sar1 Arg82 (Figure 3C) is required for recognition of Sar1 by Sec12. To test this hypothesis, we measured the nucleotide exchange activity of the D71A mutant using a quantitative biochemical reconstitution system (Futai et al., 2004; Richardson and Fromme, 2015). We monitored the kinetics of Sar1 activation by the purified Sec12 cytoplasmic domain in the presence of synthetic liposomes. The liposomes are essential in this reaction because the activation of full-length Sar1 only occurs on the surface of a membrane. We observed that Sec12ΔC[D71A] lost 93% of wild-type exchange activity (Figure 3E,F, Table S3), providing an explanation for the loss of function of this mutant *in vivo*.

In order to gain further insight into why certain interface residues appear to be more critical than others, we constructed a structural animation to visualize the conformational transitions that Sar1 undergoes during the nucleotide exchange reaction (Supplemental Movie 1). This analysis used “morphing” to highlight structural transitions between crystal structures of Sar1-GDP (Huang et al., 2001) (Figure 4A), Sar1-Sec12 (this study, Figure 4B), and Sar1-GTP (Bi et al., 2002) (Figure 4C). Importantly, morphing does not reveal the precise conformational pathway between these steps, but helps to visualize the conformational differences between each experimentally determined structural snapshot and therefore can reveal key features of the Sec12 GEF mechanism.

**Figure 4:**
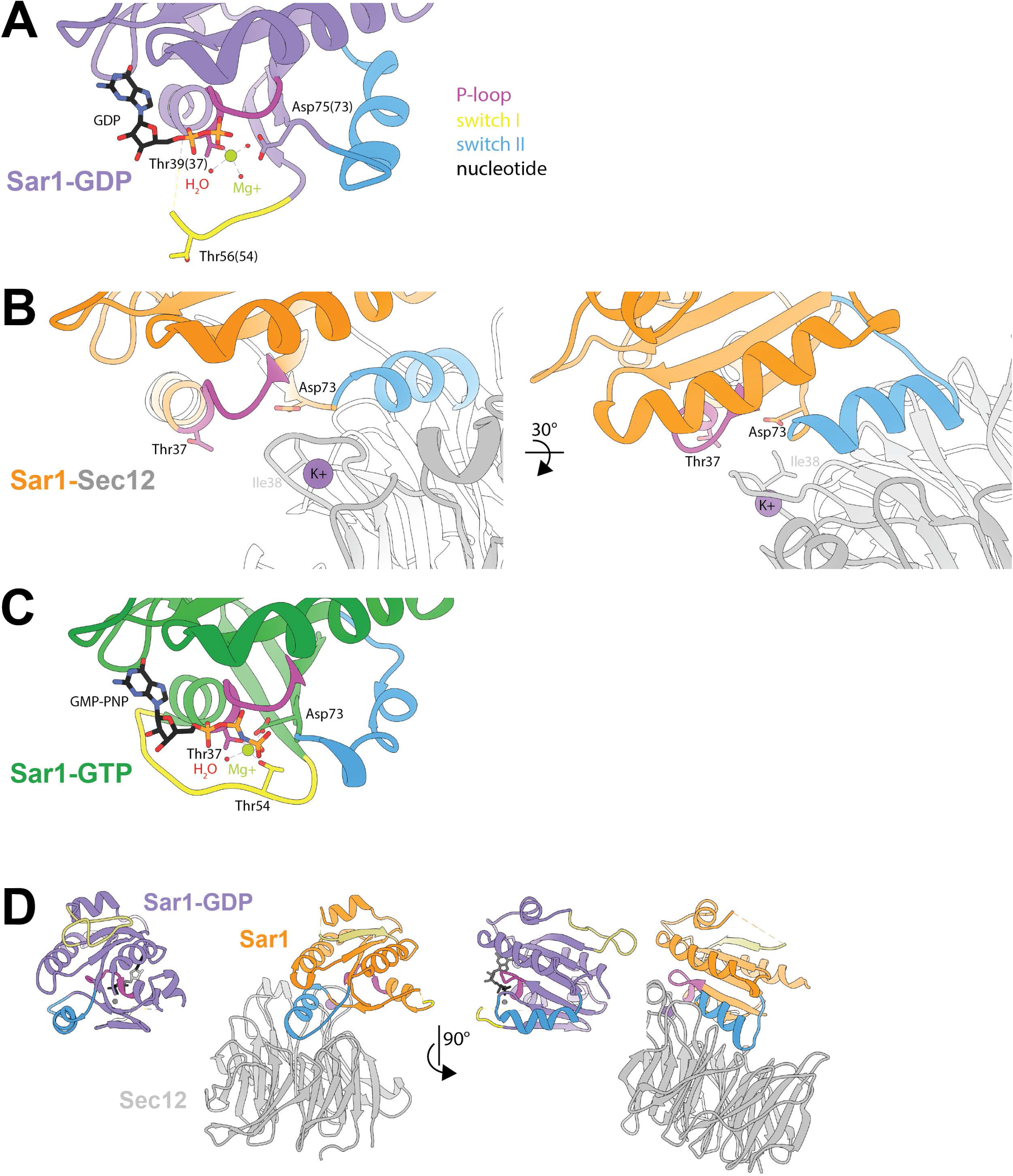
The structural basis for nucleotide release. A. Sar1-GDP (Huang et al., 2001) colored as in Figure 1, centered on the nucleotide binding pocket. Asp75 (Asp73 in yeast) is highlighted for its role in coordinating the magnesium ion for nucleotide binding. B. Sar1-Sec12 complex colored as in Figure 3, perspective as in A. Sar1 Asp73 (Asp75 in hamster) is displaced by Sec12 Ile38, disrupting a hydrogen bond network necessary for magnesium ion stabilization. C. Sar1-GTP (Bi et al., 2002) colored as in Figure 1, perspective as in A and B. D. Sar1-Sec12 complex colored as in B, next to Sar1-GDP (Huang et al., 2001) colored as in A. The Sar1 molecules are aligned, highlighting the conformational changes that occur in the switch II region as a result of Sec12 binding.

Our analysis indicates that the nucleotide release mechanism involves destabilization of nucleotide binding by steric displacement of the Sar1 switch I, switch II, and interswitch regions. The Sec12 K-loop plays an indispensable role, pushing the switch II α-helix away from the nucleotide binding site. The displaced switch II a-helix is stabilized in a new position by binding to the center of the Sec12 propeller fold (Figure 4D). Key residues mediating this interaction are Sec12 Asp71 and Tyr132 (Figure 3A,B). Sar1 switch II residue Tyr87, which in the Sar1-GDP structure is nestled on the surface of the GTPase, protrudes from Sar1 and points towards the center of the Sec12 propeller (Figure 3B). Accordingly, substitution mutations in Sec12 Tyr132 and several K-loop residues severely impacted Sec12 GEF activity (Table S3) (McMahon et al., 2012), and the importance of Sec12 Asp71 was demonstrated above.

Our structural results highlight Sec12 Ile38, which is part of the K-loop, as a critical and conserved residue for the nucleotide exchange mechanism. The Ile38 sidechain is not involved in binding to the Sec12 potassium ion (McMahon et al., 2012); instead it appears to be critical for displacing the Sar1 interswitch and switch II regions, thus repositioning Sar1 Asp73 (Figure 4B). Sar1 Asp73 is important for binding the hydrated Mg^2+^ ion and therefore stabilizing bound nucleotide, in both the Sar1-GDP and Sar1-GTP structures (Huang et al., 2001; Bi et al., 2002) (Figure 4A,C). Sec12 I38A and I38D substitution mutations disrupted nucleotide exchange (McMahon et al., 2012), likely because the mutant sidechains do not possess the bulk required to displace Sar1 Asp73 to interfere with Mg^2+^ ion binding.

### Sec12 and Sar1 interaction surfaces are distinct from structurally similar proteins

We compared the structure of the Sar1-Sec12 complex to that of the Arf1 GTPase bound to the GEF domain of the Arf-GEF Gea2 (Goldberg, 1998) (Figure 5A). Yeast Arf1 and Sar1 share ~35% amino acid sequence identity and the nucleotide-bound conformations of these proteins are structurally very similar. In contrast, the Gea2 GEF domain adopts a “Sec7” fold composed primarily of α-helices, sharing no structural resemblance with the β-propeller fold of the Sec12 cytoplasmic domain.

**Figure 5:**
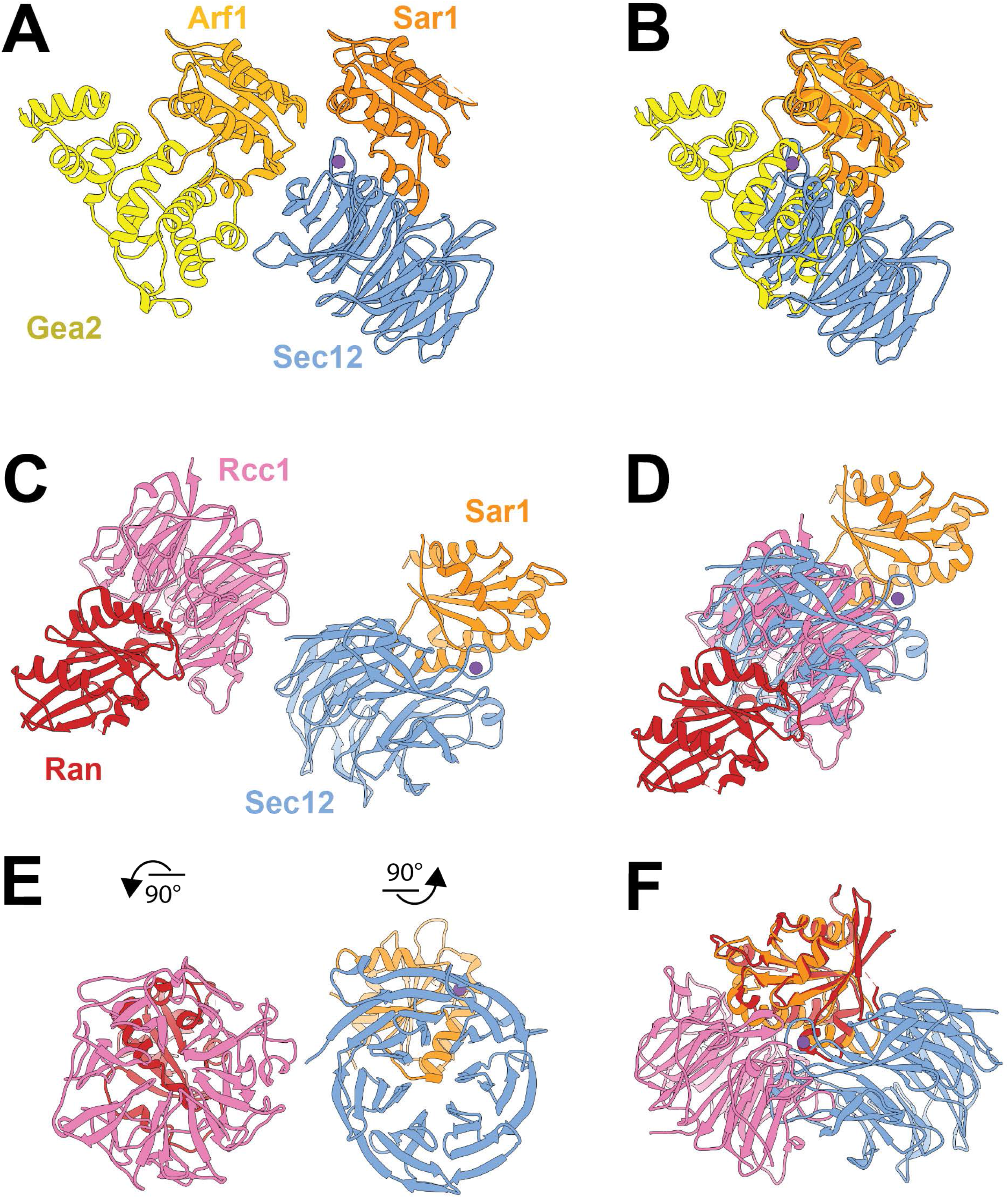
Sec12 and Sar1 interactions are distinct from structurally similar proteins. A. A ribbon representation of the structure of the Arf1-Gea2 complex (Goldberg, 1998) (Gea2 in yellow, Arf1 in gold) is shown next to the structure of Sar1-Sec12 (Sar1 in orange, Sec12 in blue). B. As in A, but with GTPases superimposed. C. A ribbon representation of the structure of the Ran-Rcc1 complex (Renault et al., 2001) (Ran in red, Rcc1 in pink) shown alongside our structure of Sar1-Sec12 (Sar1 in orange, Sec12 in blue). D. As in C, but with propeller domains superimposed. E. As in C, but the structures have been rotated in distinct 90° turns so that the GTPases lie behind the GEFs. F. As in C and D, but with the GTPases superimposed.

In spite of the similarity between Arf1 and Sar1, there are differences in their GEF-bound structures and the interfaces they use to interact with their GEFs (Figure 5B). Arf1 uses residues from both its switch I and switch II regions to interact with Gea2 (Goldberg, 1998), while Sar1 uses several residues from the phosphate loop, the interswitch region, switch II, and from helix α3.

Both Sec12 and the Sec7-domain catalyze nucleotide exchange by displacing a bound Mg^2+^ ion to weaken the bonding network of the nucleotide, but they utilize different structural approaches. Gea2 and other Sec7-domain proteins use a catalytic glutamate finger to displace the magnesium and to sterically and electrostatically repel the bound nucleotide (Beraud-Dufour et al., 1998; Goldberg, 1998). However, Sec12 does not employ an acidic residue and instead relies on a bulky hydrophobic residue (Ile38) to disrupt nucleotide and Mg^2+^ binding through steric displacement (Figure 4B).

There is one other structurally characterized GEF that adopts a β-propeller fold, Rcc1, the GEF for the Ran GTPase (Renault et al., 1998, 2001; Makde et al., 2010). Rcc1 uses a “β-wedge” protrusion from one blade of its β-propeller to interact with Ran, similar to the structure and function of the K-loop of Sec12, although the Rcc1 wedge and Sec12 K-loop lie on opposite faces of the β-propeller fold (Figure 5C,D, Figure S4). In spite of the use of comparable structural protrusions, the interactions between the GEFs and their substrate GTPases are fairly different. Ran binds to Rcc1 closer to the center of the propeller blades than does Sar1 to Sec12 (Figure 5E). Accordingly, Ran interacts with all seven blades of Rcc1, whereas Sar1 interacts with only six of the seven blades of Sec12. Both Sar1 and Ran interact with their GEFs using residues from their phosphate loop, interswitch, switch II and helix α3 regions. However, superposition of Ran and Sar1 in their GEF complex structures (Figure 5F) reveals that Rcc1 and Sec12 interact with significantly different surfaces of their GTPase substrates to achieve similar outcomes.

### Sec12 positions Sar1 for membrane insertion during activation

*In vivo,* Sec12 is anchored to the ER membrane by its C-terminal transmembrane domain. However, the orientation of the β-propeller domain with respect to the ER membrane is unknown. As activation of Sar1 coincides with the release of its amphipathic α-helix, the ability of the Sec12 cytosolic domain to orient it towards the ER membrane is important to avoid nonproductive aggregation of this helix. The cytoplasmic GEF domain appears able to directly interact with membranes *in vitro*, as it can perform membrane-dependent activation of Sar1 without its transmembrane domain (Futai et al., 2004). Only 10 residues lie between the structured portion of the cytosolic domain and the predicted transmembrane domain, suggesting that the GEF domain is constrained within a short distance of the ER membrane surface. To identify potential membrane binding surfaces, we analyzed the electrostatic surface potential of the Sar1-Sec12 complex using APBS in Chimera (Baker et al., 2001; Pettersen et al., 2004; Dolinsky et al., 2007). On one surface of the complex, we observed a prominent region of positive charge, while the remainder of the surface was predominantly negatively charged (Figure 6A). Although the residues on the positive surface are not strictly conserved, examination of homologous sequences indicates that the positively charged character of this surface of Sec12 appears to be conserved across evolution (Figure S5). Notably, the C-terminus of the Sec12 β-propeller domain, which is directly connected to the transmembrane domain by a short linker, is also located on the same face of the complex (Figure 6B), This positive surface is therefore a strong candidate for an ER membrane interaction surface. We note this surface represents an edge, rather than one of the two faces, of the β-propeller fold, and is distinct from the previously proposed membrane binding surface (McMahon et al., 2012). Interestingly, Rcc1 also uses an edge of its propeller fold to bind to the negatively charged phosphate backbone of nucleosomal DNA (Makde et al., 2010).

**Figure 6:**
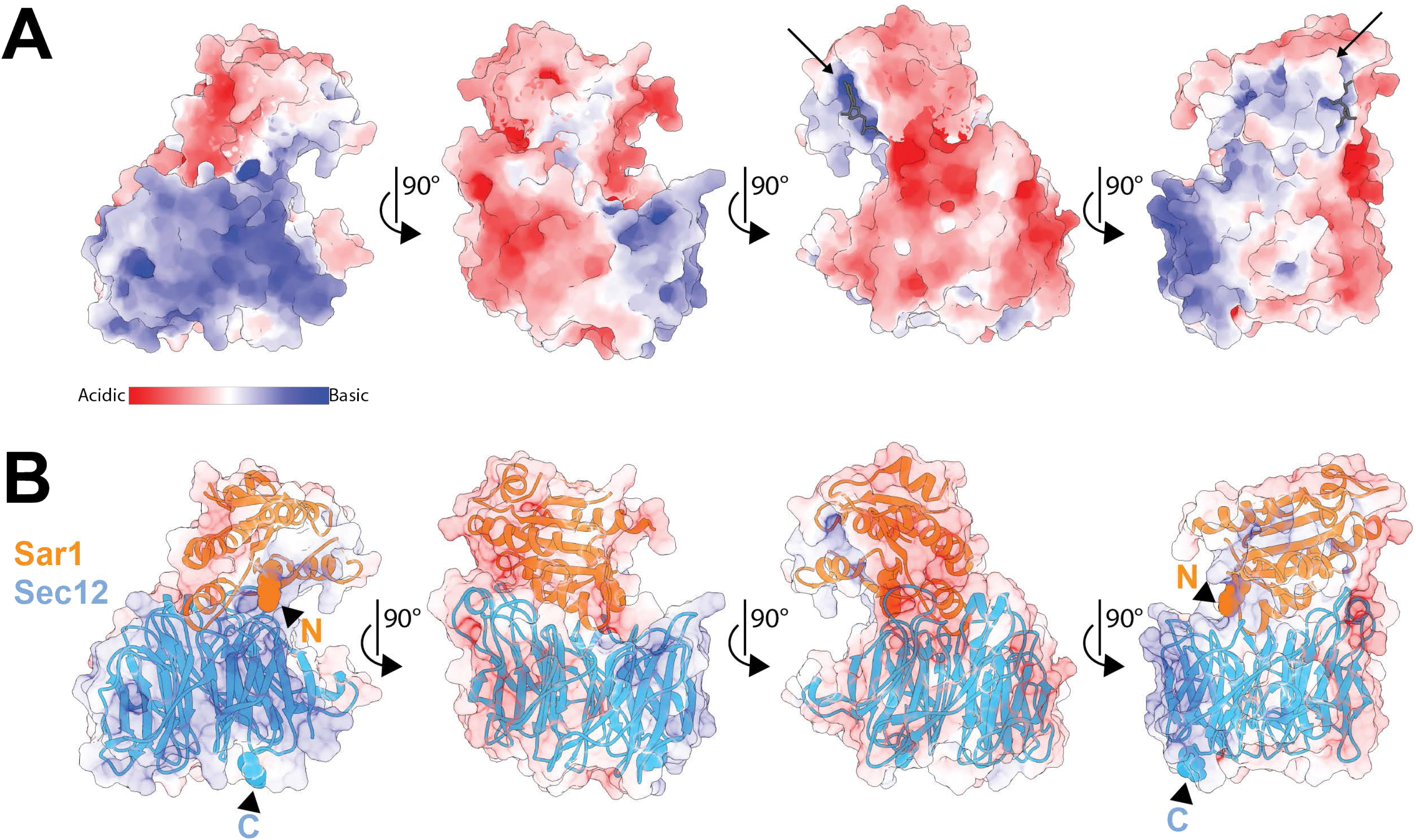
The Sec12 β-propeller domain contains a likely membrane-binding surface. A. Surface rendering of Sar1-Sec12 complex colored by electrostatic potential from −5kT/e (red) to +5kT/e (blue). A substantial positively-charged patch is observed on one face of the complex. The location of the nucleotide binding pocket is indicated with a black arrow and a nucleotide (dark gray) has been added for visualization purposes. Note that this complex is nucleotide-free. B. Similar to A, but with transparent surface and ribbon representations of Sec12 (blue) and Sar1 (orange). The N-terminal residues of the Sar1 construct (G24 and K25) are represented as orange spheres, while the C-terminal residue of the Sec12 construct (N344) is depicted as blue spheres. The termini are also indicated with arrowheads and labeled accordingly.

Upon GTP-binding to Sar1, the resulting conformational change triggers folding of the N-terminal 23 amino acids of Sar1 into an amphipathic helix that inserts into the ER membrane (Antonny et al., 1997; Huang et al., 2001). Intriguingly, Sar1 Gly24, which is the N-terminus of the Sar1 construct used for crystallization, lies on the same face of the Sar1-Sec12 complex as the positive surface and the C-terminus of the Sec12 β-propeller domain (Figure 6B and Supplemental Movie 1). Therefore, our structure leads us to a mechanistic proposal for how Sec12 can directly couple Sar1 activation to Sar1 membrane insertion (Figure 7). Due to the orientation of the Sec12 β-propeller domain on the ER membrane, when Sec12 binds to Sar1 and displaces nucleotide, it positions Sar1 so that the Sar1 N-terminus faces the membrane. Upon GTP-binding, Sar1 reveals its N-terminal amphipathic helix which inserts into the ER membrane as Sar1 dissociates from Sec12.

**Figure 7:**
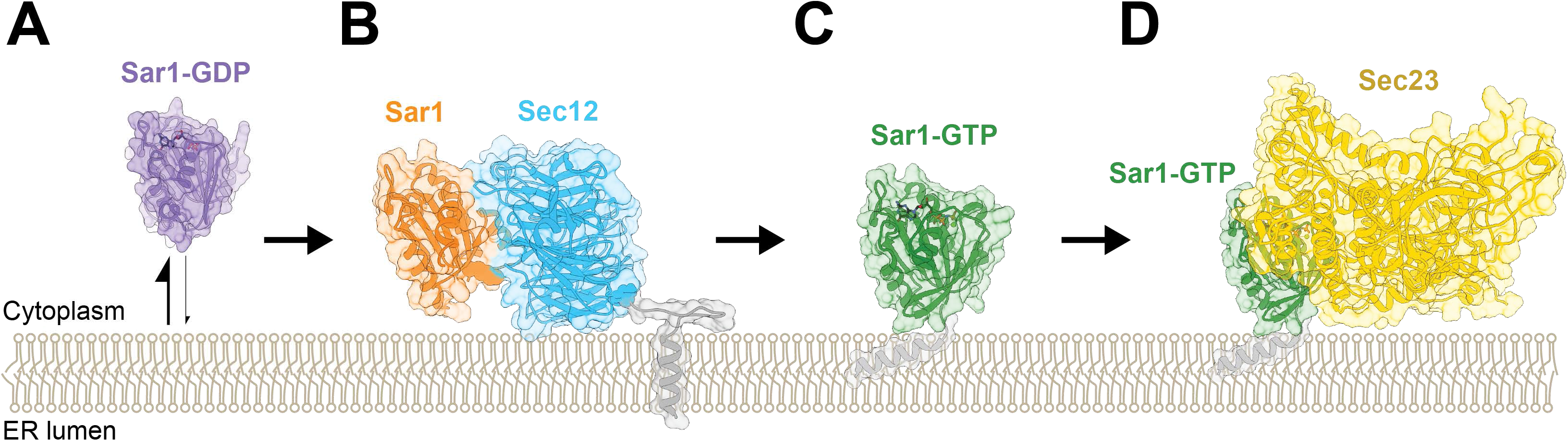
Structural model for Sar1 activation by Sec12 on the ER membrane. A. Colored in purple, Sar1-GDP (Huang et al., 2001) is cytosolic but samples the membrane surface (Antonny et al., 1997). The ER membrane is depicted as stylized lipids, approximately to scale. All proteins are shown as ribbons with a transparent surface. B. The Sar1-Sec12 complex (this study, Sar1 in orange, Sec12 in blue) interacts with the ER via its positively charged surface and the transmembrane domain of Sec12. A transmembrane domain and linker (gray) are modeled to represent how Sec12 is attached to the ER membrane. This attachment and the positively-charged patch (Figure 6) provide physical constraints for the cytosolic domain’s interaction with the membrane and position Sar1 with its N-terminus towards the membrane. The N-terminus of Sar1 and the C-terminus of Sec12 are represented as spheres. C. Membrane-bound Sar1-GTP (Bi et al., 2002) is colored in green. The N-terminal amphipathic helix is modeled in gray to indicate how Sar1 interacts with the ER membrane in its active conformation. D. As in C, but with Sec23 (yellow) (Bi et al., 2002) bound to Sar1-GTP, portraying recruitment of the first effector in COPII vesicle coat formation.

## Discussion

Activation of Sar1 by Sec12 on the ER membrane surface is the essential initiating event of the eukaryotic secretory pathway. Here we have reported the crystal structure of a Sar1-Sec12 complex. This complex represents a pivotal, nucleotide-free intermediate in the transition from the inactive (GDP-bound) to active (GTP-bound) state during GEF-mediated activation. We provide a detailed mechanistic explanation for nucleotide release and propose a mechanism for how Sec12 couples Sar1 activation with membrane insertion of its N-terminal amphipathic helix.

Although many structures of GEF-GTPase complexes have been determined, Sar1 and Sec12 serve indispensable roles to initiate the secretory pathway and possess distinguishing structural features that are vital to their function. The N-terminal amphipathic α-helix is the defining feature of Sar1 and most other Arf family members. In addition to serving as a membrane anchor, this helix has been demonstrated to serve a critical role in membrane remodeling events including induction of membrane curvature and driving membrane fission (Bielli et al., 2005; Lee et al., 2005; Drin and Antonny, 2010; Boucrot et al., 2012; Schwieger et al., 2017). Work from several laboratories has provided crucial structural insights into how Sec7-domain Arf-GEFs exchange nucleotide, and models for how Sec7-domain Arf-GEFs with PH-domains are organized on the membrane surface (Cherfils et al., 1998; Goldberg, 1998; Renault et al., 2003; DiNitto et al., 2007; Aizel et al., 2013; Malaby et al., 2013; Padovani et al., 2014; Galindo et al., 2016; Richardson et al., 2016; Karandur et al., 2017; Das et al., 2019). However, it remains unclear whether the Sec7 domain itself interacts directly with the membrane, and therefore a detailed mechanistic description of how Arf-GEFs deal with insertion of the substrate amphipathic α-helix has not been reported.

Among the known Arf-GEFs, Sec12 is unique as an integral membrane protein. Why did evolution favor a transmembrane domain GEF for Sar1 but peripheral membrane protein GEFs for other Arf family GTPases? One possibility is the location of the GEFs and GTPases. Transmembrane domain proteins localizing to the Golgi complex, endosomes, and plasma membrane are initially synthesized at the ER before trafficking to their destination organelle. Perhaps the transient localization of a plasma membrane or endosomal Arf-GEF at the ER would result in deleterious activation of Arf substrates ectopically at the ER.

We took advantage of the constraint provided by the Sec12 transmembrane domain, together with a striking positively charged region, to identify the likely membrane-interacting surface of the Sec12 β-propeller GEF domain. When bound to Sec12, Sar1 is oriented such that its N-terminus is pointed towards the membrane. Upon GTP binding to Sar1, Sec12 will dissociate while the Sar1 N-terminal amphipathic α-helix takes shape in just the right place to insert into the membrane. We think the orientation of the complex on the membrane is important because optimal positioning would greatly increase the frequency of productive encounters between the folding amphipathic α-helix and the membrane surface. This would bias outcomes towards successful insertion events rather than aggregation-prone exposure of the hydrophobic surface of this helix to the cytoplasm.

We observed an additional interesting feature regarding the surface electrostatics of the Sar1-Sec12 complex. The Sar1 nucleotide binding pocket is a small positively charged “bullseye” within the large negatively charged surface of the Sar1-Sec12 complex that faces away from the membrane (Figure 6A, arrows). Because nucleotides bear a strong negative charge due to their phosphate groups, we suspect the membrane-distal surface acts as an electrostatic “funnel” (Carpenter and Lightstone, 2016) to guide GTP to its binding pocket in Sar1.

It has been known for some time that cellular K^+^ levels diminish in response to metabolic stress (Findlay, 1994) and depletion of cellular K^+^ to ~25% of normal levels inhibits ER-Golgi transport, likely by blocking exit from the ER (Judah et al., 1989). Sec12 is a tantalizing candidate for the focal point of this regulation of trafficking. The potassium ion-binding K-loop is essential for Sec12 function in cells (McMahon et al., 2012), and our structure demonstrates how the K-loop acts to directly disrupt the Sar1 nucleotide-binding pocket by displacing the switch and interswitch regions.

## Materials and Methods

### Protein purification

For purification of full-length Sar1, we used a previously reported plasmid (McMahon et al., 2012) using the pET21b (Novagen) backbone that expresses full-length Sar1 from *Saccharomyces cerevisiae* with a C-terminal hexahistidine tag. For crystallization, we mutated this plasmid to create an N-terminal truncation of the first 23 amino acids (corresponding to the amphipathic helix) of Sar1 (ΔN23-Sar1). Purification of both protein constructs was similar: plasmid was transformed into the Rosetta2 strain of *E.coli* (Novagen). Two liters of culture were grown in terrific broth for 8-12h at 37°C (until the OD_600_ ~2). The temperature was then reduced to 16°C and 1 hour later IPTG was added to a final concentration of 300μM to induce protein expression. Following overnight expression (14-18h), cells were collected via centrifugation and resuspended in lysis buffer (40mM Tris pH 8, 300mM NaCl, 10% glycerol, 10mM imidazole, 1x PMSF, and 1mM DTT). Cells were lysed by sonication and ΔN23-Sar1-6xHis was purified by nickel affinity chromatography (Ni-NTA resin, Qiagen) and eluted with elution buffer (40mM Tris pH 8, 300mM NaCl, 10% glycerol, 250mM imidazole, and 1mM DTT). The sample was further purified using anion exchange chromatography with a linear gradient (buffer A: 20mM Tris pH 8, 1mM DTT; buffer B: 20mM Tris pH 8, 1M NaCl, 1mM DTT) on a MonoQ column (GE Healthcare). The peak fractions were collected, concentrated, and frozen in liquid nitrogen prior to storage at −80°C.

A similar approach was used to purify the cytoplasmic domain of Sec12. We used a previously reported plasmid (McMahon et al., 2012) with the pET21b (Novagen) backbone that expresses a C-terminal truncation of Sec12 (Sec12ΔC, residues 1-354, lacking the transmembrane domain and a short ER lumenal domain) from *Saccharomyces cerevisiae* with a C-terminal hexahistidine tag. Mutations were introduced via a two-step PCR protocol. For all Sec12ΔC constructs, the purification protocol used was identical to that of Sar1, except 200mM KCl was added to the nickel purification buffers and KCl was added to the MonoQ fractions to a final concentration of ~130mM.

For nucleotide exchange assays, proteins were further purified via gel filtration chromatography on a Superdex 200 Increase 10/300 column (GE Healthcare) prior to usage. Buffer for full-length Sar1: 10mM Tris pH 8.0, 150mM NaCl, 1mM DTT; buffer for Sec12ΔC constructs: 10mM Tris pH 8.0, 100mM NaCl, 100mM KCl, 1mM DTT (Note: two overlapping peaks were observed for Sec12 and both were combined). Proteins were aliquoted, flash frozen, and stored at −80°C. Prior to use, aliquots were thawed, and pre-spun for 3 minutes at ~20,000g.

### Sar1-Sec12 Complex formation

Purified Sec12ΔC and ΔN23-Sar1 (previously frozen fractions from the anionic exchange step) were mixed in a 1:3 molar ratio and incubated overnight at room temperature with calf intestinal alkaline phosphatase (Sigma). After 12-18h, the sample mixture was subjected to size exclusion chromatography on a Superdex 200 Increase 10/300 column (GE Healthcare) in 10mM Tris pH 7, 250mM KCl, 1mM DTT. The trace displayed two dominant peaks corresponding to ΔN23-Sar1:Sec12ΔC complex and free ΔN23-Sar1 (Figure S1). The fractions containing the complex were pooled and concentrated to ~34mg/mL in a 10kDa molecular weight cutoff concentrator from Millipore.

### Crystallization

An Art-Robbins Phoenix robot was used to set up 96-well crystallization screens containing several concentrations of purified ΔN23-Sar1:Sec12ΔC at room temperature and 4°C using the sitting drop vapor diffusion method. Several initial crystallization hits were optimized but found to contain only Sec12. Diffraction data used for Sar1-Sec12 structure determination was collected from a single crystal. This crystal grew from a 5mg/mL protein solution mixed with 200mM ammonium sulfate, 100mM MES pH 6.5, 30% PEG 5000 MME after 3 weeks at 4°C. The crystal was washed in place with a solution of 50% glycerol as cryoprotectant then flash frozen in liquid nitrogen.

### Diffraction data collection, structure solution, model building, and refinement

X-ray diffraction data were collected on NE-CAT Beamline 21-IDE using an Eiger 16-M detector. X-ray data were processed by the RAPD pipeline (http://necat.chem.cornell.edu/) which utilizes XDS (Kabsch, 2010). The structure was solved by molecular replacement using Phaser (McCoy et al., 2007) in Phenix (Liebschner et al., 2019), using previously published structures of yeast Sec12 (PDB:4H5I) (McMahon et al., 2012) and hamster Sar1-GDP (PDB:1F6B) (Huang et al., 2001) as search models. Iterative rounds of model building in Coot (Emsley et al., 2010) and refinement in Phenix were performed to yield a final model with reasonable statistics (Table 1).

### Nucleotide exchange assays

GEF assays were performed essentially as described previously (Futai et al., 2004; Richardson and Fromme, 2015). Intrinsic tryptophan fluorescence of Sar1 was measured in real time using a fluorometer (Photon Technology International). Assays were carried out under the following conditions: 2μM Sar1, 10nM Sec12ΔC construct, 200μM GTP, 333μM major-minor liposomes (Futai et al., 2004) in HKM buffer (20mM HEPES pH 7.4, 160mM KoAc, 5mM MgCl_2_) for 10 minutes at 30°C.

All assays were performed in triplicate. The raw traces were cropped to contain just the exchange section. These traces were fit by a single exponential using Prism 8 (GraphPad). The rate constant (k) of the intrinsic Sar1 exchange was subtracted from all other rate constants. The resulting rate constant was divided by the [GEF] to calculate the exchange rate (k_exchange_). The average of 3 exchange rates was plotted along with the 95% confidence interval. For trace visualization, one representative trace is shown after being normalized to between 0 and 1.

### Yeast complementation assays

Cell viability was maintained in a *sec12Δ::KANMX* haploid background strain (CFY4048) by the presence of a SEC12:URA3 centromeric plasmid. A *SEC12-3xHA*:LEU2 centromeric plasmid was subjected to site-directed mutagenesis and then transformed into CFY4048. Cells were plated in a dilution series onto -LEU and 5-FOA plates and grown for 2 days at 30°C and 37°C before being imaged. Nonfunctional *sec12* alleles are unable to support cell growth.

### Surface analyses

Conservation analysis was carried out using the Consurf web server at default settings (Armon et al., 2001; Landau et al., 2005; Ashkenazy et al., 2010; Celniker et al., 2013; Ashkenazy et al., 2016). The electrostatic surface potential was calculated using the APBS plugin for Chimera (Pettersen et al., 2004) and the map was colored on a red (−5kT/e) to blue (+5kT/e) scale (Baker et al., 2001; Dolinsky et al., 2007), after charges were added to the structure using the Add Charges feature in Chimera (using the AMBER ff14SB setting for standard residues and AM1-BCC for others).

### Software

All software used for the above analyses is maintained in our laboratory by SBGrid (Morin et al., 2013). Molecular graphics created with UCSF Chimera and ChimeraX (Pettersen et al., 2004; Goddard et al., 2018). The animated video, depicting transitions between Sar1 conformations in three different crystal structures (GDP-bound, nucleotide-free, and GTP-bound), was generated using the “Morph” feature in Pymol (Schrödinger, LLC). Note that the conformational pathway generated by morphing does not necessarily represent the actual pathway of the GEF reaction, but is helpful for highlighting the structural differences between the states observed in the crystal structures.

### Sequence alignment

The multiple sequence alignment shown in Figure S5 was performed using Clustal Omega with default settings (Madeira et al., 2019).

### Data deposition

The X-ray diffraction structure factors and coordinates of the model have been deposited in the RCSB PDB (Accession code: 6X90).

## Acknowledgments

We thank Fred Hughson for sharing plasmid reagents. We thank Ian Price for assistance with data collection and processing. We thank Saket Bagde, Steve Halaby, and Brian Richardson for critiquing the manuscript. This work is based upon research conducted at the Northeastern Collaborative Access Team beamlines, which are funded by the National Institute of General Medical Sciences from the National Institutes of Health (P30 GM124165). The Eiger 16M detector on 24-ID-E is funded by a NIH-ORIP HEI grant (S10OD021527). This research used resources of the Advanced Photon Source, a U.S. Department of Energy (DOE) Office of Science User Facility operated for the DOE Office of Science by Argonne National Laboratory under Contract No. DE-AC02-06CH11357. This work was supported by National Institutes of Health grants R01GM098621 and R35GM136258 to J.C.F., by an Alfred P. Sloan Foundation Fellowship and a National Science Foundation Graduate Research Fellowship (grant DGE-1144153) to A.M.N.J. Any opinions, findings, conclusions, or recommendations expressed in this material are those of the authors and do not necessarily reflect the views of the funders.

## Author contributions

AMNJ --conceptualization and design, data acquisition/investigation, data analysis/interpretation, writing and revising; JCF --conceptualization and design, data analysis/interpretation, writing and revising, funding acquisition.

## Supplemental Materials

**Supplemental Movie 1:**

Inactive Sar1 is depicted in orange with bound GDP (yellow) and a magnesium ion (green). The view zooms in to focus on the bound magnesium ion, which is coordinated through a hydrogen bond network that includes Sar1 residue Asp75 (yeast residue Asp73). While the distance between Asp75 and the magnesium ion is measured at 3.8 Å, the Asp75 actually directly hydrogen bonds with a water molecule which in turn hydrogen bonds with the magnesium ion (water molecules are not shown). After zooming out, Sec12 approaches and binds to Sar1-GDP. The K loop directly displaces Asp75, disrupting the hydrogen bond network that holds the magnesium ion in place. As the magnesium ion leaves, Sar1 undergoes significant structural rearrangements in its switch I and interswitch regions, while the switch II region moves as a rigid body towards the center of Sec12. The structural rearrangements and the destabilization from the absence of the magnesium ion cause GDP to dissociate. After zooming out again, the entire Sar1-Sec12 structure is visible. The view centers on residue Ile38 of Sec12 and then displays the 3.2 Å measured distance between it and residue Asp73 of Sar1 (hamster residue Asp75). This distance is appropriate for these residues which are in close contact but do not clash. This is in contrast to the hypothetical distance measured between the GDP/GTP-bound Sar1 conformations which would be approximately 0.5-1.5 Å (not shown). The view zooms out to reveal the entire Sar1-Sec12 structure and rotates 90°. Spheres appear to denote the C-terminal residue of Sec12 (residue 344, which is adjacent to its TMD) and the N-terminal residue of Sar1 (residue 24, which is close to the amphipathic helix). The complex rotates back 90° to show the orientation that it is likely to adopt on the ER membrane, with both termini residing on the same face directed downwards. Then, the view centers on the nucleotide binding pocket of Sar1 as GTP moves into position. Upon GTP binding, structural rearrangements in Sar1 occur and Sec12 dissociates. Ultimately, nucleotide binding is stabilized by a magnesium ion which is coordinated through a hydrogen bond network involving Asp73 of Sar1. In the GTP-bound conformation, the switch I region of Sar1 is structured and visible as a loop at the forefront of the frame. Sar1-GDP (PDB:1F6B) and Sar1-GTP (PDB:1M2O) were the starting and ending models, respectively (Huang et al., 2001; Bi et al., 2002)

**Figure S1:**
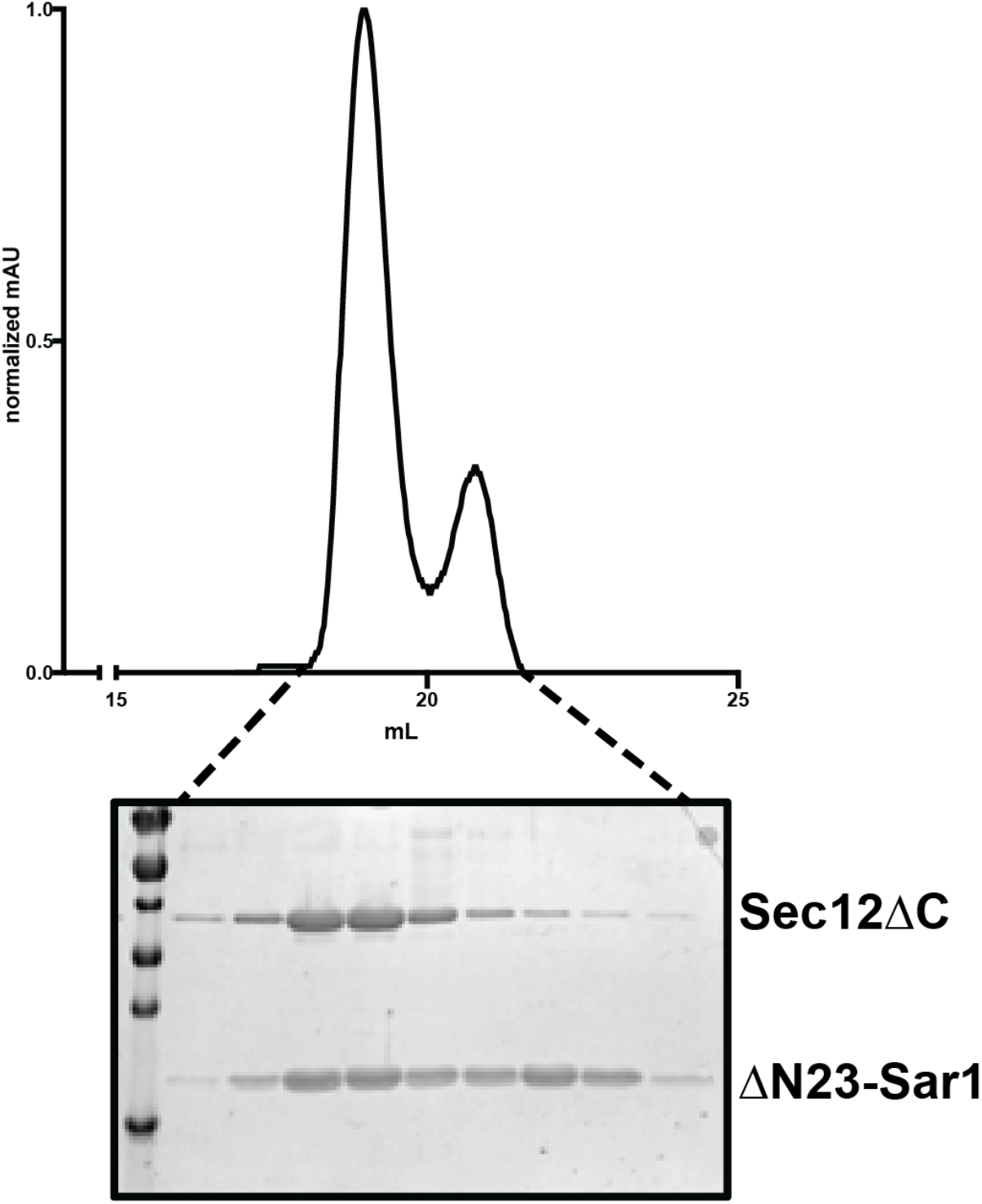
Sec12-Sar1 complex formation. Top, Trace of UV absorption during size-exclusion chromatography. The first peak corresponds to Sar1 in complex with Sec12, while the second peak represents unbound Sar1. Bottom, A Coomassie-stained SDS-PAGE gel showing the peak fractions.

**Figure S2:**
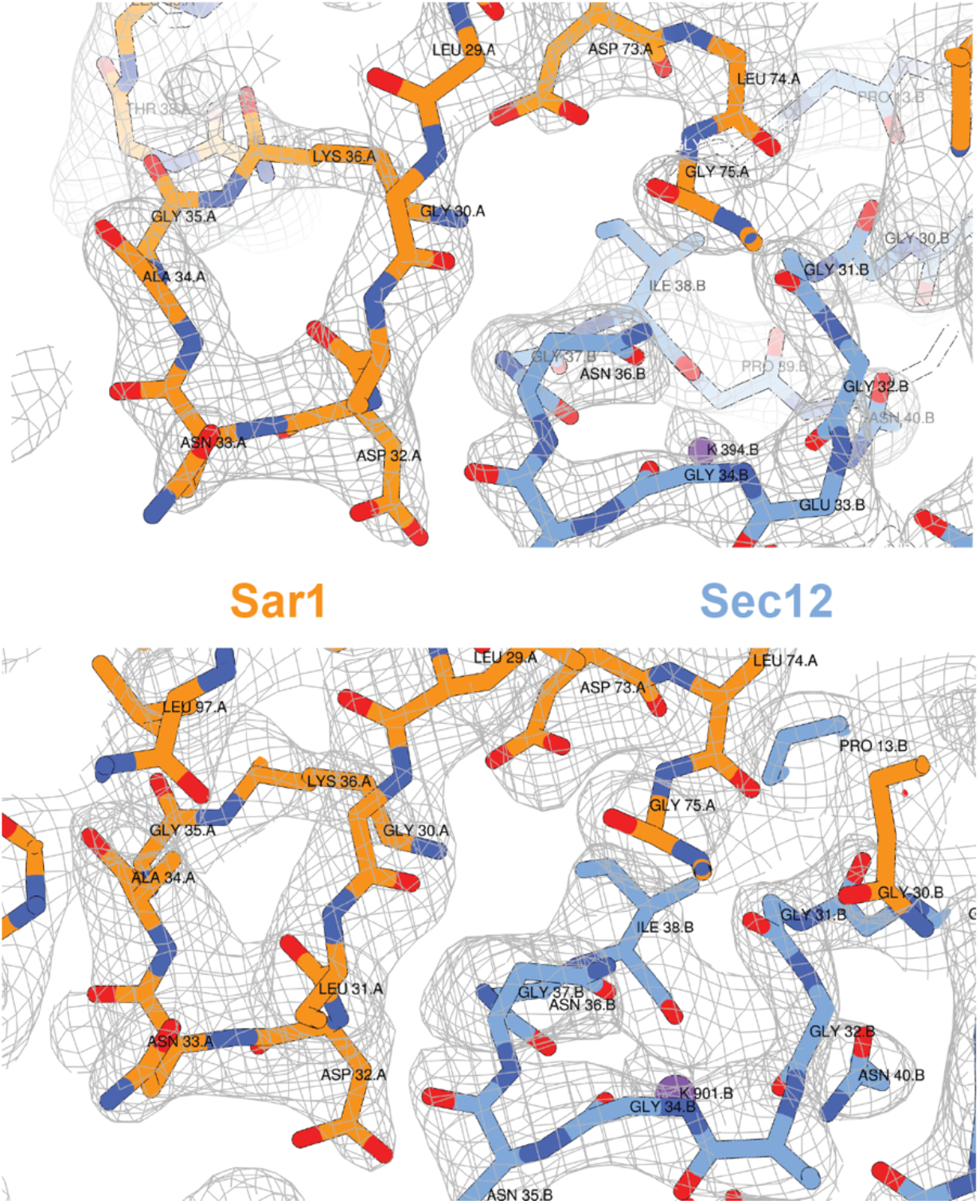
Example electron density maps. Top, Weighted 2Fo-Fc electron density map, displayed at 1.5σ, in the region of the complex comprising the interface between Sar1 (orange) and the Sec12 K-loop (blue). Bottom, Simulated annealing composite omit map, displayed at 1.0σ, in the same region.

**Figure S3:**
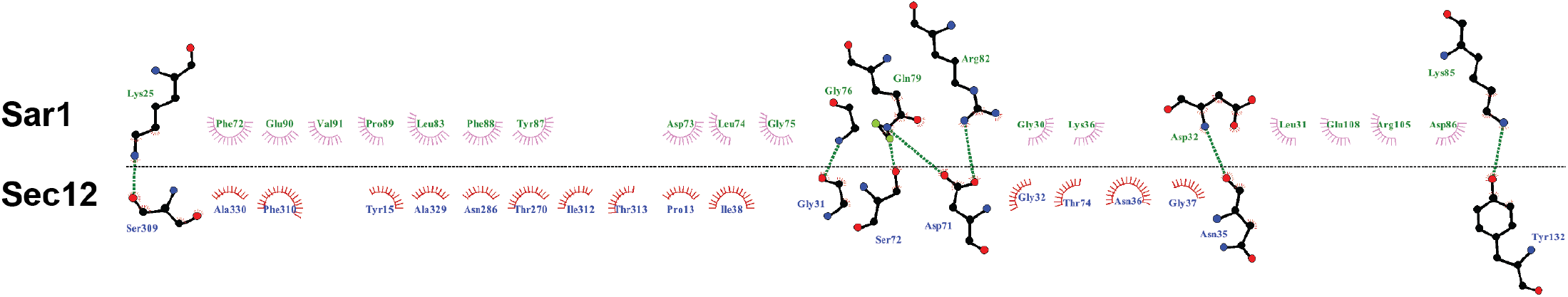
2D-depiction of the Sar1-Sec12 interaction interface. Schematic 2D representation of the interface between Sar1 and Sec12. Residues from Sar1 are on top with green labels. Residues from Sec12 are below with blue labels. Residues involved in hydrogen bonding are shown as stick representations with the hydrogen bond indicated by a dashed green line. Residues with hydrophobic interactions across the interface are depicted as labeled semi-circles with outward lines. This diagram was generated with LigPlot+ (Laskowski and Swindells, 2011).

**Figure S4:**
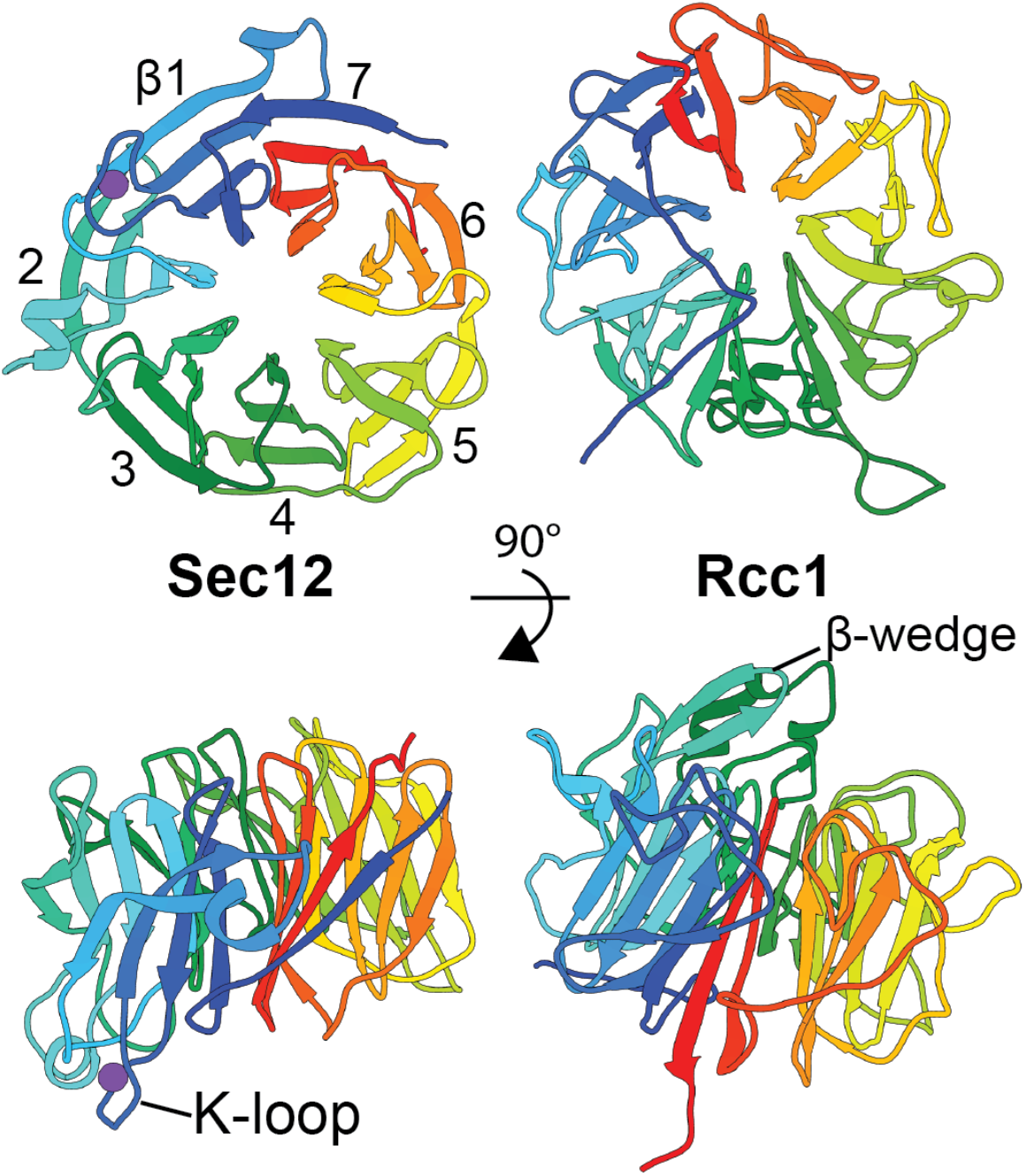
Comparison of Sec12 and Rcc1 β-propeller folds. Top, Sec12 (left) and Rcc1 (right) are aligned then colored from N-(blue) to C-terminus (red). Each contains seven blades that form a propeller. The blades of Sec12 are labeled from 1-7. The first N-terminal β-strand of Sec12 forms a β-sheet with the three final C-terminal β-strands of the cytoplasmic domain to form the seventh blade. However, the first two N-terminal β-strands of Rcc1 join with the two final C-terminal β-strands to form the seventh blade (Renault et al., 1998, 2001; Makde et al., 2010). Bottom, Rotated 90° to show that the functional protrusions (Sec12: K-loop, Rcc1: β-wedge) are on opposite faces.

**Figure S5:**
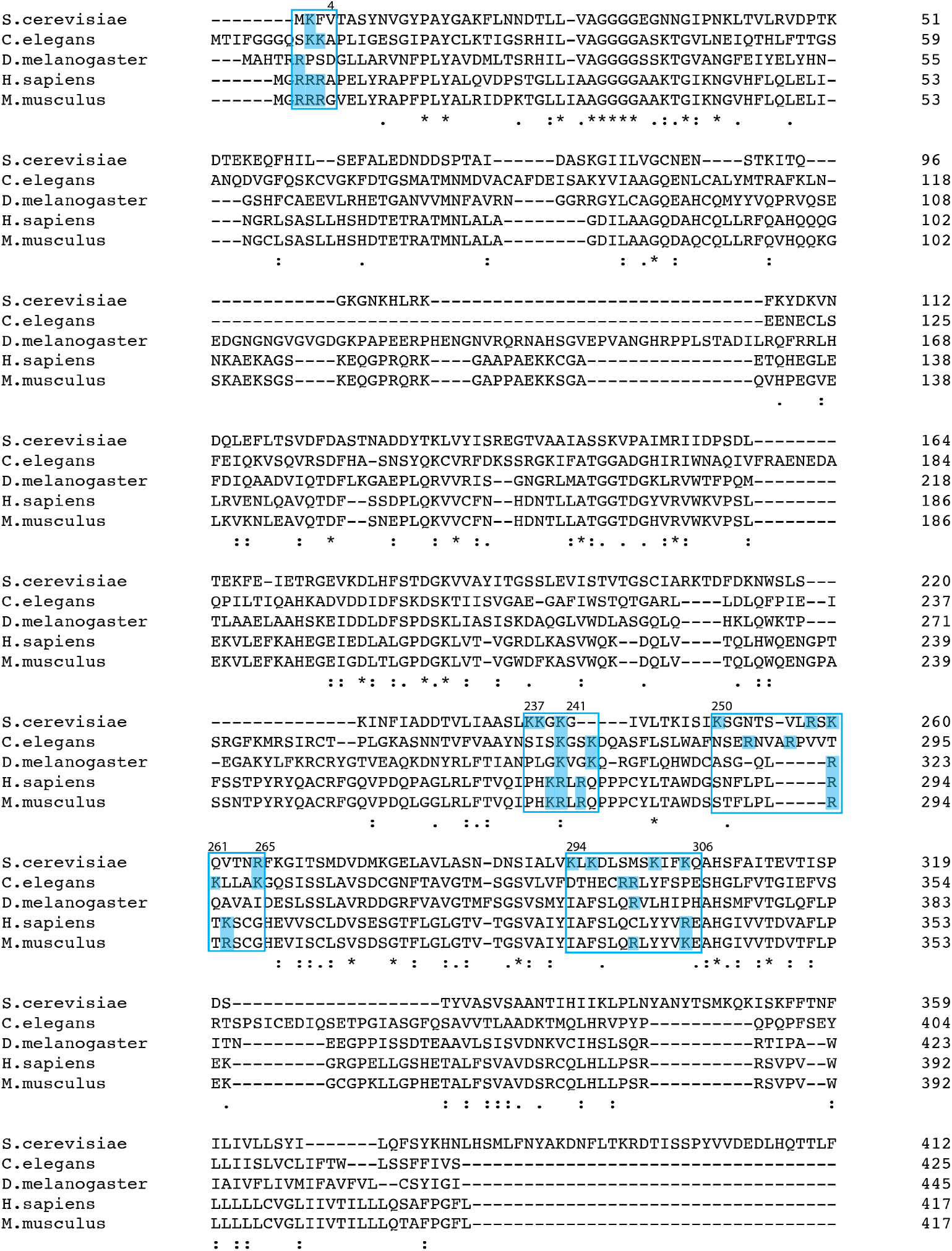
Sec12 sequence alignment highlighting positively-charged surface. A sequence alignment of Sec12 homologues from several species created with Clustal Omega (Madeira et al., 2019). The residues corresponding to the proposed membrane-proximal surface are enclosed in blue boxes. The basic residues within those regions are highlighted with a blue background. The total number of positive charges on this surface range from 5 in *D. melanogaster*, 10 in *C. elegans*, 12 in *S. cerevisae* and *H. sapiens*, and 13 in *M. musculus*.

**Table S1:** Diffraction data and refined model statistics.

|  |  |
| --- | --- |
| Wavelength | 0.979180 |
| Resolution range | 57.71 – 2.258 (2.339 – 2.258) |
| Space group | P 21 21 21 |
| Unit cell | 50.333 93.62 115.423 90 90 90 |
| Total reflections | 174650 (16809) |
| Unique reflections | 26271 (2524) |
| Multiplicity | 6.6 (6.5) |
| Completeness (%) | 99.06 (97.07) |
| Mean I/sigma (I) | 20.01 (1.20) |
| Wilson B-factor | 66.3 |
| R-merge | 0.05526 (1.278) |
| R-meas | 0.05999 (1.388) |
| R-pim | 0.02305 (0.5334) |
| CC1/2 | 0.999 (0.844) |
| CC* | 1 (0.957) |
| Reflections used in refinement | 26101 (2514) |
| Reflections used for R-free | 1222 (122) |
| R-work | 0.2557 (0.4923) |
| R-free | 0.3051 (0.5320) |
| CC(work) | 0.944 (0.815) |
| CC(free) | 0.910 (0.768) |
| Number of non-hydrogen atoms | 3756 |
| Macromolecules | 3753 |
| Ligands | 1 |
| Solvent | 2 |
| Protein residues | 486 |
| RMS(bonds) | 0.004 |
| RMS(angles) | 0.97 |
| Ramachandran favored (%) | 94.13 |
| Ramachandran allowed (%) | 5.87 |
| Ramachandran outliers (%) | 0 |
| Rotamer outliers (%) | 0.48 |
| Clashscore | 4.11 |
| Average B-factor | 95.59 |
| Macromolecules | 95.6 |
| Ligands | 93.6 |
| Solvent | 72.5 |

**Table S2:**
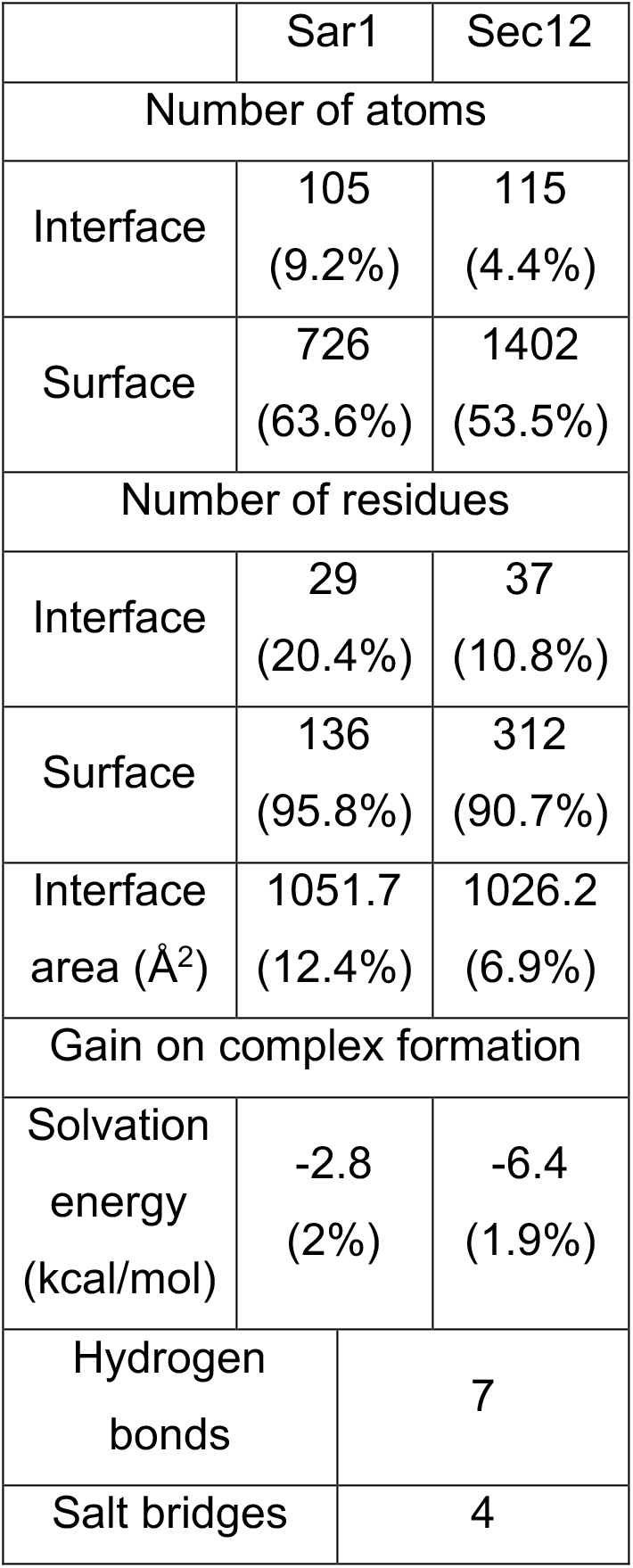
Analysis of Sar1-Sec12 interaction surface.

|  | Sar1 | Sec12 |
| --- | --- | --- |
| Number of atoms |  |  |
| Interface | 105<br>(9.2%) | 115<br>(4.4%) |
| Surface | 726<br>(63.6%) | 1402<br>(53.5%) |
| Number of residues |  |  |
| Interface | 29<br>(20.4%) | 37<br>(10.8%) |
| Surface | 136<br>(95.8%) | 312<br>(90.7%) |
| Interface<br>area (Å<sup>2</sup>) | 1051.7<br>(12.4%) | 1026.2<br>(6.9%) |
| Gain on complex formation |  |  |
| Solvation<br>energy<br>(kcal/mol) | -2.8<br>(2%) | -6.4<br>(1.9%) |
| Hydrogen<br>bonds | 7 |  |
| Salt bridges | 4 |  |

**Table S3:**
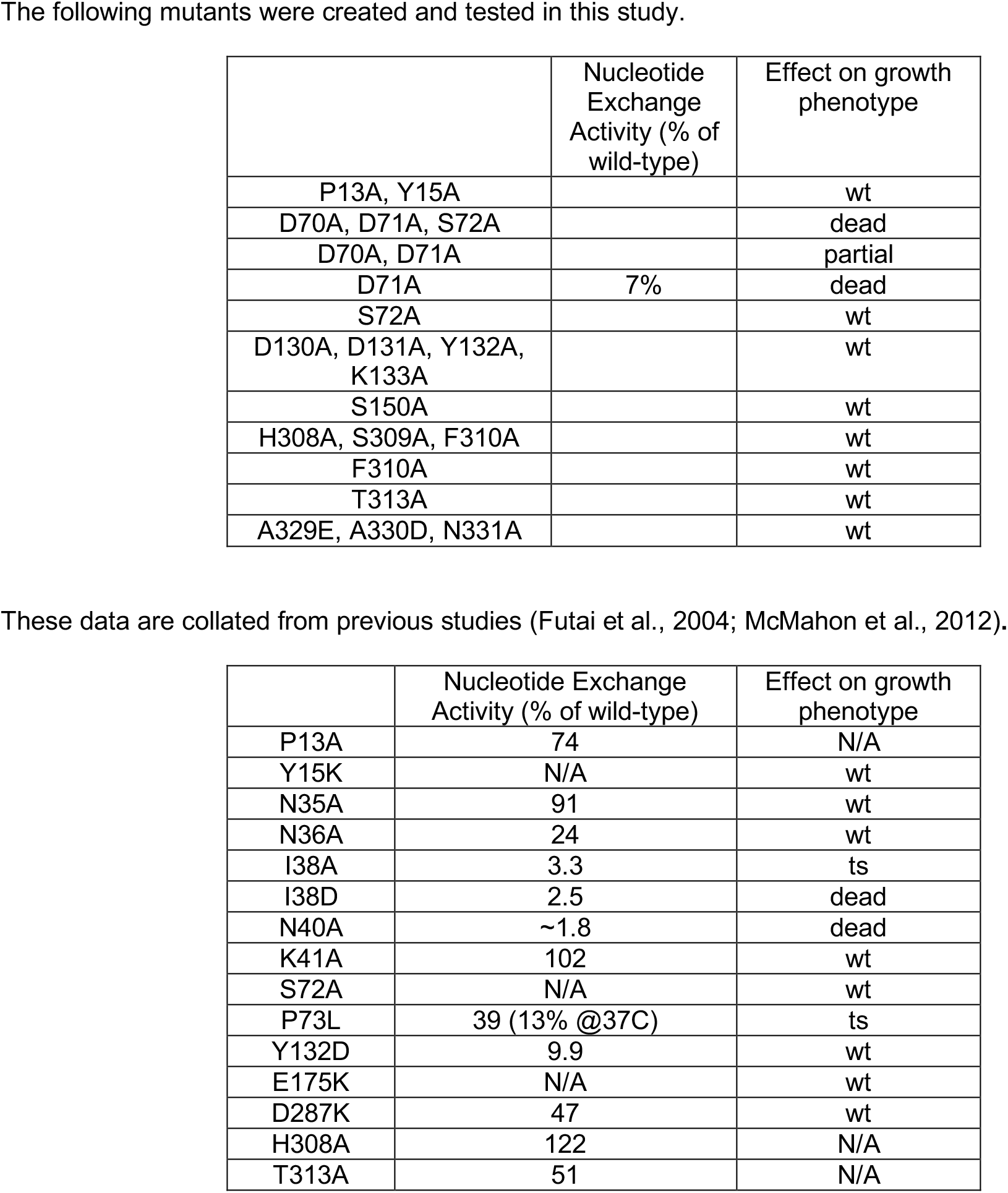
Phenotypes of Sec12 mutations.

|  | Nucleotide Exchange Activity (% of wild-type) | Effect on growth phenotype |
| --- | --- | --- |
| P13A, Y15A |  | wt |
| D70A, D71A, S72A |  | dead |
| D70A, D71A |  | partial |
| D71A | 7% | dead |
| S72A |  | wt |
| D130A, D131A, Y132A, K133A |  | wt |
| S150A |  | wt |
| H308A, S309A, F310A |  | wt |
| F310A |  | wt |
| T313A |  | wt |
| A329E, A330D, N331A |  | wt |

|  | Nucleotide Exchange Activity (% of wild-type) | Effect on growth phenotype |
| --- | --- | --- |
| P13A | 74 | N/A |
| Y15K | N/A | wt |
| N35A | 91 | wt |
| N36A | 24 | wt |
| I38A | 3.3 | ts |
| I38D | 2.5 | dead |
| N40A | ~1.8 | dead |
| K41A | 102 | wt |
| S72A | N/A | wt |
| P73L | 39 (13% @37C) | ts |
| Y132D | 9.9 | wt |
| E175K | N/A | wt |
| D287K | 47 | wt |
| H308A | 122 | N/A |
| T313A | 51 | N/A |

